# The macroecology of viral coinfection

**DOI:** 10.1101/2025.09.08.674542

**Authors:** Cecilia A. Sánchez, Colin J. Carlson, Amy R. Sweeny

## Abstract

Coinfection is common in wild animals, and can profoundly influence disease outcomes and transmission. However, most coinfection research is based on laboratory experiments or a few well-studied wildlife systems. Here, we use data from the PREDICT project – the largest standardized wildlife disease surveillance project ever conducted – to describe patterns of viral coinfection and evaluate their association with host and virus factors. Within the viruses prioritized for testing, we find that coinfection is rare (detected in just 223 of 65,662 animals), but still more common than expected by chance – especially coinfections of coronaviruses, paramyxoviruses, and influenza A virus. We further find that coinfection is associated with host age, but bats and rodents exhibit opposing relationships. We find that captive wild animals (specifically rats and mallards) exhibit higher coinfection than free-ranging wild animals, highlighting potential risks to wildlife and human health from the wildlife trade. Finally, we find that biases in sampling and testing can shape observed patterns of coinfection. Our results characterize associations among viruses in the largest relevant dataset currently available. Factors associated with coinfection in this dataset such as host-level variation, virus-virus interactions, and human interference should be further tested via surveillance and laboratory approaches to better resolve the role of coinfection as an important and likely overlooked driver of viral dynamics in nature.

## Introduction

Parasites (including microparasites and macroparasites) exist in complex communities, where coinfection with multiple parasites at any time is the norm for wild animals (Cox 2001; Petney and Andrews 1998). Coinfections provide an opportunity for direct interactions between parasites, such as competition for resources (Budischak et al. 2017) or space (Knowles et al. 2013), or indirect interactions, such as those mediated by host immune responses (Ezenwa et al. 2021). These interactions can have a dramatic effect on outcomes at the host level, and transmission dynamics at the population level (Telfer et al. 2010; Gorsich et al. 2018; Pedersen and Fenton 2007; Graham 2008).

Several theoretical frameworks in disease ecology have incorporated the dynamics and consequences of coinfection (Pedersen and Fenton 2007; Fenton and Pedersen 2005; Graham 2008; Sweeny et al. 2021), but it can be difficult to test their predictions against empirical evidence. Within-host processes are easiest to study in laboratory settings (Fenton et al. 2014), but the controlled settings that make experimental approaches useful (e.g., laboratory organisms from a single host species, with known ancestry and no unknown parasites) also limit their similarity to real-world dynamics in natural communities. A few well-developed (and often long-term) field systems have also led to a number of insights about interactions within diverse host and parasite communities (Ezenwa et al. 2021), and some systems allow for natural experiments such as parasite removal or nutritional supplementation (Sweeny et al. 2020; Knowles et al. 2013; Ezenwa and Jolles 2015). However, neither field nor experimental case studies can provide insight into the full network of interactions occurring within parasite communities (or their consequences) at the ecosystem scale or above.

Whereas large-scale biodiversity data have reached volumes that allow for unprecedented research and monitoring, wildlife disease data collection has historically been fragmented between emerging infectious disease surveillance, wildlife management efforts, and ecological research. Across all three fields, there are significant global gaps in surveillance (Carlson et al. 2025; Cohen et al. 2023), and – despite broader progress towards open science in ecology and epidemiology – the sharing of spatially-explicit, host-level infection data remains rare (Schwantes et al. 2025; Albery et al. 2022). Some studies have used data synthesis to develop global infection datasets (Cohen et al. 2023, 2020; Heckley and Becker 2024; Warmuth et al. 2023), but in most cases, they have focused on single-host, single-parasite relationships, rather than single-host, multi-parasite dynamics.

Here, we explore coinfection dynamics at the global scale for the first time, using the single largest standardized study of wildlife disease ever conducted. The PREDICT project was a decade-long, multi-country initiative funded by the U.S. Agency for International Development (USAID) that conducted viral surveillance primarily in wild animals (PREDICT Consortium 2020). Focused on virus families of public health concern but capturing a sizable fraction of wildlife virus diversity, the project produced one of the largest standardized multi-virus datasets ever assembled and resulted in the discovery of new viruses (Anthony et al. 2017b; Goldstein et al. 2018), an improved understanding of drivers of virus infection and diversity (Anthony et al. 2017a; Eskew et al. 2025), and insight into human and virus-related risk factors for zoonotic spillover (Saylors et al. 2021). However, minimal work has been done using the PREDICT data to test macroecological hypotheses. Using these data, we set out to explore three questions: (1) How common is viral coinfection? (2) Is coinfection more common in certain host groups, life stages, and settings? And finally: (3) Are there non-random patterns of coinfection driven by specific virus taxa, or by associations between specific pairs of virus taxa? Through this work, we also aimed to identify promising avenues for future research on viral coinfection.

## Methods

### Data cleaning

Analyses were performed using the R statistical environment v4.5.1 (R Core Team 2025). All steps of the analysis pipeline (data cleaning, statistical modeling, figure generation) can be reproduced using the code available at https://github.com/viralemergence/macroecology-coinfection. Data from the PREDICT-2 project (2015-2019; the second half of the decade-long initiative) were openly shared on the USAID Development Data Library prior to the takedown of the USAID website. Our analyses relied on two PREDICT datasets: an animal sampling dataset containing details such as species, sex, and age class and a second dataset containing virus test results for individual specimens (which also included some basic host information). We cleaned the test result dataset by excluding human results and fixing minor inconsistencies in animal host taxonomy (e.g. host scientific name was mostly provided at the species level, but sometimes only at the genus or family level). We modified the animal taxonomic groups originally provided by joining “birds” and “poultry/other fowl” into one “birds” group, and separating a “rodents/shrews” group into “rodents” and “shrews” groups. Two detections of the hantavirus Thottapalayam virus were originally classified as belonging to the Bunyaviridae family; we reclassified them to the Hantaviridae family to align with current virus taxonomy(Kuhn et al. 2024). Finally, we joined the two datasets so that each test result was paired with the relevant animal data.

### Data overview

Specimen collection (*n* = 126,952 unique specimens from 65,662 unique animals) was concentrated in countries in West and Central Africa as well as South and Southeast Asia, with > 10,000 specimens collected in Sierra Leone, Guinea, and Bangladesh each (Fig. S1). Most specimens were collected from bats (*n* = 77,334, 60.9%) and rodents (*n* = 25,683, 20.2%). The dataset included some farm and companion animals, including poultry, dogs, and swine. Virus screening was conducted using consensus polymerase chain reaction (PCR) assays (714,954 total tests), mostly targeting virus families of significance to human health (e.g., coronaviruses, filoviruses, and paramyxoviruses), but in some cases testing for specific viruses of interest (e.g., influenza viruses and ebolaviruses). Further details on PREDICT can be found in the PREDICT Legacy Book (https://ohi.vetmed.ucdavis.edu/programs-projects/predict-project/reports).

### Analyses

Across all animals and within each animal taxonomic group (16 total groups; Table S1), we calculated infection prevalence (number infected divided by number tested) and coinfection prevalence (number coinfected divided by number tested). We considered an animal to be infected when one or more test results for specimens collected from that animal were positive for at least one virus, and to be coinfected when test results were positive for two or more unique viruses. For bats, rodents, and birds (the three largest taxonomic groups), we compared the observed coinfection prevalence to the expected pairwise prevalence based on the prevalence of each virus, using a permutation testing approach. Specifically, we randomized infection across individual animals while maintaining the observed prevalence of each virus within the taxonomic group, then calculated the number of simulated coinfected animals. We repeated this procedure 1000 times for each taxonomic group, then plotted the simulated number of coinfected animals compared to the number observed in the PREDICT data.

To explore patterns of coinfection among virus taxa, we created a coinfection matrix at the virus family level, where each count within the matrix represents a coinfection of two virus families occurring in the same animal. We included virus family coinfections that represented detections of two viruses in the same family (e.g. two coronaviruses) or in different families (e.g. one coronavirus and one paramyxovirus). For animals infected with three viruses, we counted each coinfection separately (e.g., an animal infected with coronavirus X, coronavirus Y, and paramyxovirus Z would count for two coronavirus-paramyxovirus coinfections and one coronavirus-coronavirus coinfection). We used the *ggraph* package (Pedersen 2024) to visualize the frequency of virus family coinfection. To test if certain virus families were co-detected more often than expected by chance, we first calculated the expected coinfection prevalence of two virus families as the product of each virus family’s overall prevalence in the dataset. We then performed a binomial test for each virus family–virus family combination to determine if the observed coinfection prevalence was significantly greater than expected. To account for multiple comparisons, we adjusted *P* values using a Bonferroni correction. Finally, we repeated the above analyses for two subsets of the full data: bats and rodents. We also created a coinfection matrix at the individual virus level and visually explored coinfection patterns; however, due to high sparsity in this matrix, we did not analyze these data further.

For each infected animal *(n* = 3,271), we quantified the number of unique viruses detected in its test results and classified its coinfection status as either single infection (exactly one virus detected) or coinfection (two or more viruses detected). We then used generalized linear models to explore correlates of coinfection status. For all model datasets, we excluded taxonomic groups with ≤ 10 animals (carnivores, cattle/buffalo, dogs, goats/sheep, other), animals that were missing sex or age class data, and neonates and fetuses (*n* = 809 total animals excluded). **1)** For an all-animals model (*n* = 2,462), we used the ‘glm’ function in the *stats* package (R Core Team 2025) to model animal coinfection status as a function of geographic region (Africa and West Asia; South, East, and Southeast Asia), taxonomic group (bats, birds, rodents, shrews), sex (female, male), age class (adult, subadult, juvenile), captivity status (wild free-ranging, owned domesticated, wild in captivity), and the number of virus families tested for across all specimens associated with that animal. Given inconsistency in host identification (i.e., not all animals were identified to species level), we did not include any species-level traits as model predictors. **2)** For a bats-only model *(n* = 1,519), we classified whether each bat species is known to roost in caves using data from Willoughby *et al*. (Willoughby et al. 2017) and the IUCN Red List. We excluded bats for which cave roosting behavior was not available and bats not identified to at least family level. Using the ‘glmmTMB’ function in the *glmmTMB* package (Brooks et al. 2017), we modeled coinfection status as a function of five fixed effects variables (geographic region, sex, age class, cave roosting status, number of virus families tested), and one random effects variable (host family). **3)** For a rodents-only model (n = 488), we modeled coinfection status as a function of sex, age class, and number of virus families tested. Subadults and juveniles were collapsed into one age class category due to sample size restrictions, while host family and geographic region were not included as variables because nearly all rodents were murids and were collected in South, East, and Southeast Asia.

## Results

We found that both infection and coinfection were rare overall for viruses prioritised in sampling. Out of 65,662 unique animals, 3,271 (4.98%) were infected with one or more viruses (Fig. 1A); virus prevalence in birds (487/5,617; 8.67%) and bats (2,059/36,181; 5.69%) was somewhat higher than average, while prevalence in rodents (549/12,876; 4.26%) was slightly lower than average (Table S1). Animals with at least one virus detection were most often captured in South, East, and Southeast Asia (Fig. S2). Within infected animals, there was uneven geographic distribution of animal taxonomic groups; infected bats, rodents, and shrews were widely distributed, while infected birds and swine were found only in South, East, and Southeast Asia (Fig. 1B).

**Figure 1.**
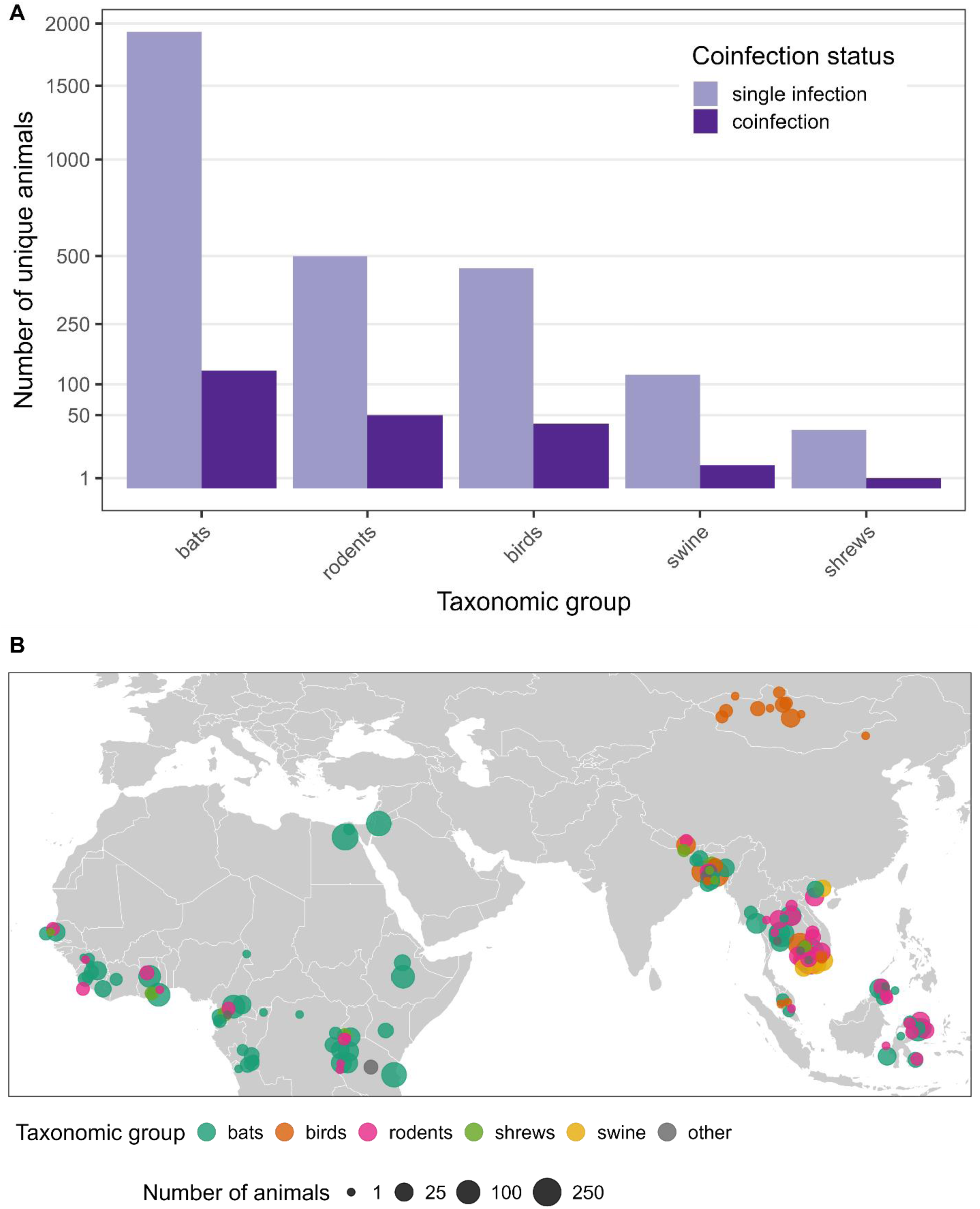
**A.** Number of unique animals infected with one or more viruses, separated by taxonomic group. Groups in which only single infections were observed are not displayed (dogs: *n* = 10; goats/sheep: *n* = 4; cattle/buffalo: *n* = 2; carnivores: *n* = 1; other: *n* = 1). **B.** Distribution of animals infected with one or more viruses, colored by taxonomic group. Point size corresponds to the number of animals captured at a given location; coordinates were truncated so that nearby locations could be grouped together. Points are partially transparent and jittered for improved visibility.

A single virus was detected in most infected animals (*n* = 3,048, 93.18%); coinfections of two (*n* = 211, 6.45%) or three (*n* = 12, 0.37%) viruses in the same animal were infrequent. Of the 223 coinfected animals, over half were bats (*n* = 128, 57.4%) with the remainder being 50 rodents, 39 birds, 5 swine, and 1 shrew (Table S1). Within infected animals, coinfection rates were highest in rodents (50/549, 9.11%), birds (39/487, 8.01%), and bats (128/2,059, 6.22%). Roughly half of the triple coinfections (7/12; Table S2) were domesticated ducks and geese, which were all infected with duck coronavirus, influenza A virus, and either Newcastle virus or avian paramyxovirus 6; the remainder were bats infected with a mix of coronaviruses, paramyxoviruses, influenza A virus, and a novel rhabdovirus.

### Coinfection is non-random across phylogeny and life stage

Bats, rodents, and birds exhibited unique patterns of coinfection, even accounting for their over-representation in the dataset. Coinfections were more common than expected based on the observed prevalence of all viruses: for all three taxonomic groups, we applied a permutation testing approach, and found that observed coinfection rates exceeded the maximum simulated coinfection rates (across 1,000 simulations for each taxonomic group; Fig. S3). However, when accounting for geographic region, sex, age class, captivity status, and number of virus families tested in a generalized linear model, we found that both birds (odds ratio (OR) = 0.24, 95% confidence interval (CI): 0.09-0.54, *p* = 0.001) and rodents (OR = 0.55, 95% CI: 0.33-0.89, *p* = 0.018) were significantly less likely to be coinfected than bats (Fig. 2A, Table S3). These findings could support the widespread hypothesis (and growing evidence) that bats are able to tolerate viral infection, and potentially coinfection, through a unique set of immune adaptations (Letko et al. 2020; Irving et al. 2021; Becker et al. 2025). However, we note that coinfection rates within the all-animals model dataset (i.e., animals with available sex and age data) differed compared to the coinfection rates in the dataset of all 3,048 infected animals: while coinfection was still high in rodents (46/488, 9.43%) and bats (112/1,628, 6.88%), it was noticeably lower in birds (12/316, 3.80%).

**Figure 2.**
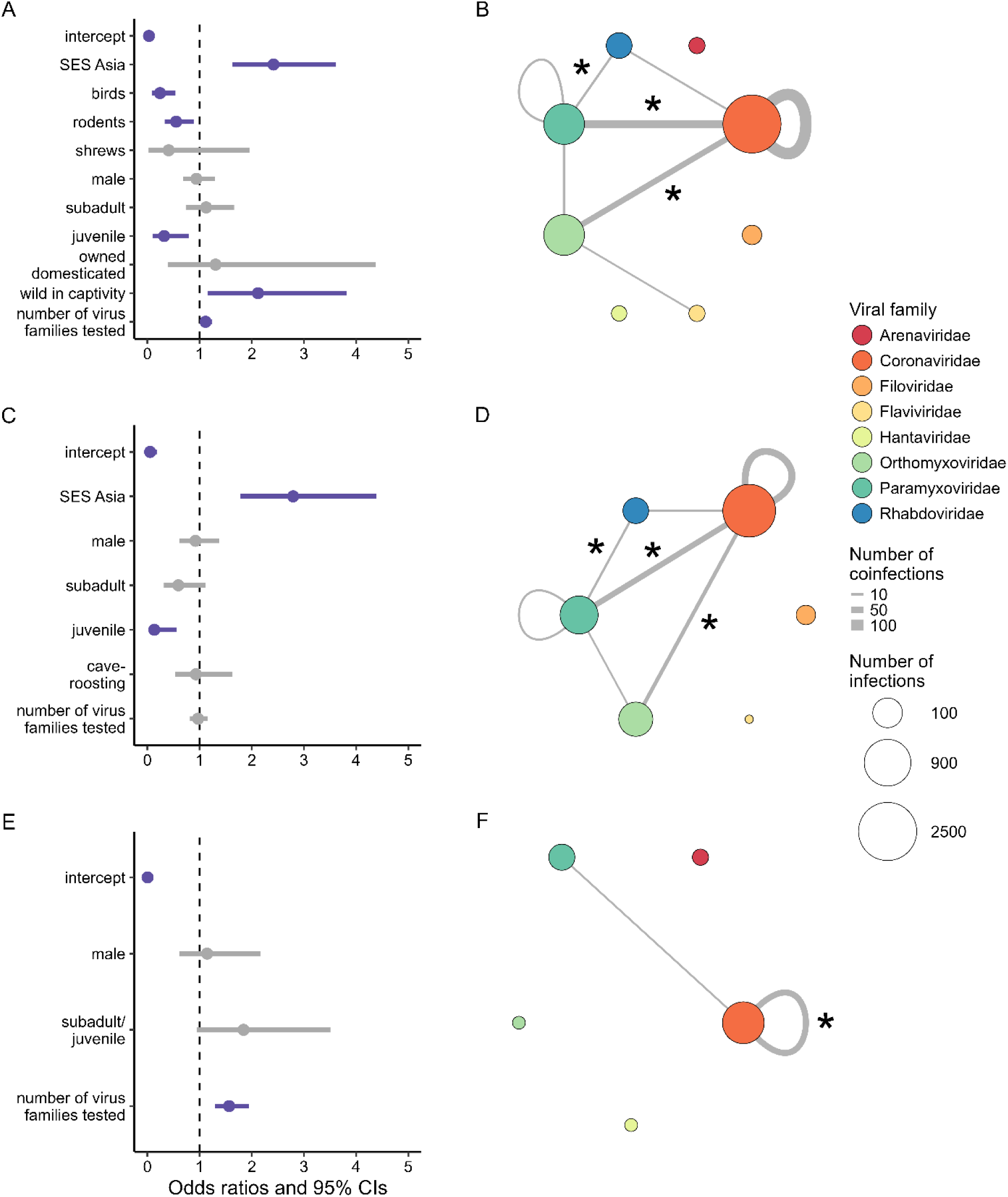
Predictors of virus coinfection (**A**, **C**, **E**) and virus family coinfection networks (**B**, **D**, **E**). **Left column:** generalized linear model outputs (odds ratios and 95% confidence intervals (CIs)) for predictors of coinfection are shown for all animals (**A**), bats only (**C**), and rodents only (E). CIs that do not cross the vertical line at 1 (i.e., statistically significant) are purple. Reference levels for categorical predictors are Africa and West Asia (geographic region), bats (taxonomic group), female (sex), adult (age class), wild free-ranging (captivity status), and non-cave roosting (roosting status). SES Asia = South, East, and Southeast Asia. **Right column:** networks of virus family coinfection are shown for all animals (**B**), bats only (**D**), and rodents only (**F**). Nodes represent virus families, while edges represent the frequency of virus family coinfection. Asterisks indicate coinfections that occurred significantly more than expected.

Coinfection was associated with life history, but those relationships varied between groups. Based on previous studies (Anthony et al. 2017a; Peel et al. 2025), we hypothesized that infection prevalence – and therefore coinfection rates – would be higher in younger animals, due to waning of maternal antibodies and less-developed immune systems (Goenka and Kollmann 2015). We found that younger rodents (juveniles and subadults) were more likely to be coinfected than adults, though this result was not statistically significant (OR = 1.84, 95% CI: 0.95-3.51, *p* = 0.066; Fig. 2E, Table S5). However, juvenile bats were less likely to be coinfected than adults (OR = 0.13, 95% CI: 0.03-0.55, *p* = 0.006; Fig. 2C, Table S4), and this effect was strong enough that it dominated the all-animals model (OR = 0.32, 95% CI: 0.10-0.80, *p* = 0.030; Fig. 2A, Table S3). This pattern is aligned with the widespread hypothesis that bats’ unique immune adaptations allow them to tolerate persistent viral infections (Calisher et al. 2006; Plowright et al. 2016), which might accumulate over the course of an individual’s lifespan (Aguillon et al. 2025). Alternatively, depending on the timing of sampling by the PREDICT study, juvenile bats may have been captured at times when levels of maternal antibodies were still high. Though logistically challenging, longitudinal sampling within populations can help elucidate patterns of virus infection and coinfection in relation to host maturation (Wacharapluesadee et al. 2018; Brook et al. 2019; Peel et al. 2025).

Finally, we found that host social structure could drive coinfection patterns, but found divergent effects between natural and un-natural settings. For bats, cave-roosting behavior was not a significant predictor of coinfection status (OR = 0.93, 95% CI: 0.53-1.63, *p* = 0.796; Fig. 2C, Table S4). Cave-roosting bats can exist in large groups in close proximity, potentially increasing infection rates, but other roosting strategies can also create large, close aggregations of bats (e.g., tree-roosting *Pteropus* species that cluster together). Finer-scale contact patterns that we could not examine here (e.g., segregation by age or sex, or centrality in social networks) might lead to more correlated infection risk across viruses at the individual level. Sickness behaviors (e.g., reduced social interactions) after an initial infection could also reduce an individual’s subsequent likelihood of infection with other viruses (Ripperger et al. 2020; Moreno et al. 2021).

In contrast, across all animals, we found that wild animals in captivity were significantly more likely to be coinfected than free-ranging wild animals (OR = 2.12, 95% CI: 1.16-3.82, *p* = 0.013; Fig. 2A, Table S3). These results support the hypothesis that captivity can increase parasite transmission through both crowding, stress, and unusual patterns of interspecific contact. This hypothesis has been strongly supported by previous analyses of coronavirus testing conducted by the PREDICT project (Anthony et al. 2017b; Huong et al. 2020), and is a particular point of concern, given the potential for coinfection to lead to recombinant coronaviruses (Wells et al. 2023), and the high risk of spillover in wildlife farms and markets (Sánchez et al. 2021). However, we note that of the 163 animals classified as “wild animals in captivity” in the all-animals model dataset, 131 (80.4%) were rats (*Rattus* spp.) and 28 (17.2%) were mallards (*Anas platyrhynchos*). Further work quantifying coinfection rates in captive counterparts of a broader range of species is needed to better understand if these results are generalisable, and to examine the role of potential mechanisms in driving coinfection in captivity.

### Sampling and testing biases can shape observed coinfection patterns

Animals sampled in South, East, and Southeast Asia were more likely to be coinfected than those sampled in Africa and West Asia in both the all-animals model (OR = 2.42, 95% CI: 1.63-3.61, *p* < 0.001; Fig. 2A, Table S3) and the bats-only model (OR = 2.79, 95% CI: 1.78-4.39, *p* < 0.001; Fig. 2C, Table S4). Rather than reflecting any characteristic of one region versus another, this result might be driven by sampling artifacts. For instance, flying foxes (*Pteropus* spp.) had a high coinfection rate (all-animals model dataset: 37/245; 15.1%) and were sampled only in South, East, and Southeast Asia. Rats (*Rattus* spp.) were sampled in both geographic regions, but infected animals were almost entirely found in South, East, and Southeast Asia, and coinfection was high within this group (44/403; 10.9%). Virus family testing effort was also positively correlated with coinfection in the all-animals model (OR = 1.11, 95% CI: 1.00-1.24, *p* = 0.042; Fig. 2A, Table S3) and the rodents model (OR = 1.57, 95% CI: 1.29-1.94, *p* < 0.001; Fig. 2E, Table S5).

### Evidence of virus-virus associations

Coinfection networks revealed that certain viruses–namely, coronaviruses (CoVs), paramyxoviruses (PMVs), and orthomyxoviruses (specifically, influenza A virus)–were most frequently detected together (Fig. 2B, 2D, 2F). Notably, there were 108 CoV-CoV coinfections (57 in bats, 48 in rodents), 69 CoV-PMV coinfections (44 in bats, 2 in rodents), 45 CoV-influenza coinfections (23 in bats), and 10 PMV-influenza coinfections (1 in bats). Binomial tests indicated that CoV-PMV, CoV-influenza, and PMV-rhabdovirus coinfections occurred significantly more than expected in all animals and in bats, while coinfections with multiple coronaviruses occurred significantly more than expected in rodents (Tables S6-S8). All other virus family coinfections occurred < 10 times, and no coinfections involving arenaviruses, filoviruses, or hantaviruses were detected.

Examining coinfection patterns at the level of individual viruses, we found that viruses that are transmitted by poultry and other livestock (influenza A virus, Newcastle disease virus, and infectious bronchitis virus) had the highest degree centrality in the network (Fig. 3). However, a handful of commonly occurring betacoronaviruses, including murine coronavirus, PREDICT CoV-16 and PREDICT CoV-17, and Bat coronavirus HKU9, were also well connected within the network.

**Figure 3.**
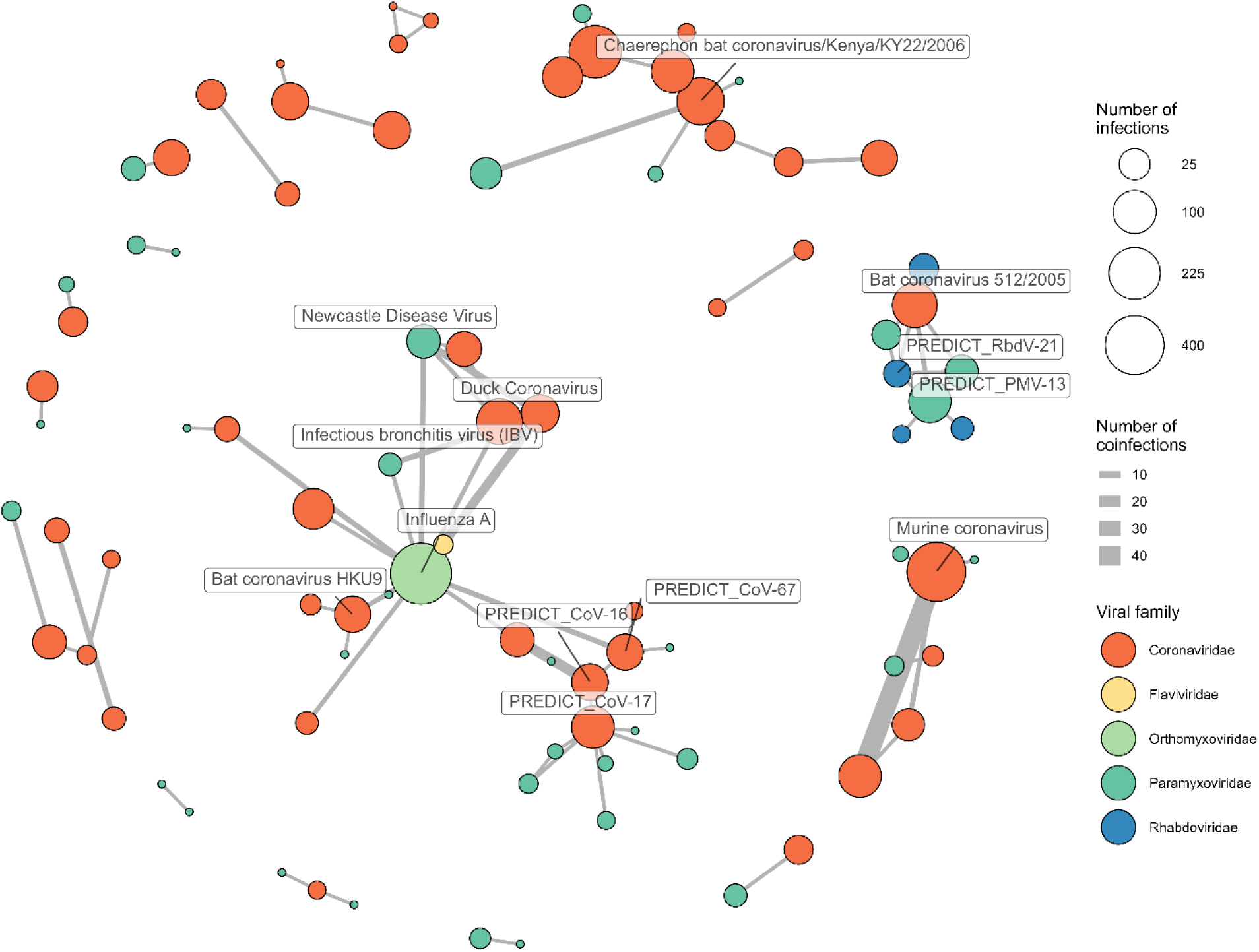
Virus coinfection network. Nodes represent individual viruses, while edges represent the frequency of virus coinfection. Only nodes with degree centrality ≥ 4 are labeled, and nodes not involved in coinfections are not displayed.

## Discussion

Here, we conducted the first analysis of viral coinfection patterns across multiple continents and host taxonomic groups. We found that within the virus groups sampled, coinfection is rare – detected in just 223 of 65,662 animals – but occurs more frequently than expected, particularly among specific virus groups. Our findings suggest that widely-occurring viruses could be important to account for when researching other pathogens. For example, influenza A virus was often detected in combination with other viruses; one possible explanation for this finding is that influenza A virus might alter host susceptibility to secondary infections. Future work could explore how specific viral coinfections influence factors such as the magnitude and duration of shedding, particularly as it could relate to zoonotic spillover risk (Schuh et al. 2025). We also found coinfection dynamics that suggest further testing would be beneficial to resolve whether these associations are modulated by broad differences in host immunology and/or fine-scale host contact processes, including in anthropogenic settings like farms and the wildlife trade. These findings are also of potential importance for public health, as they highlight the need for both basic research on host-parasite interactions, and surveillance of environments where coinfection can produce new reassortant and recombinant viruses with epidemic or pandemic potential (Anthony et al. 2017a).

Although our study is the largest macroecological study of coinfection to date, our findings should be interpreted in light of the limitations of our data. Most importantly, the PREDICT project was not designed with ecological research as the primary objective, and so the testing data have uneven coverage both across and within taxonomic groups. For instance, the PREDICT project was almost single-handedly responsible for bats becoming the best-studied wildlife hosts of viruses (Gibb et al. 2022). Given that over 60% of specimens were collected from bats, these data probably drove many of our broad-scale findings. However, only half of all bat families (10 of 21) were sampled, and an emerging body of work highlights that while bats are unique viral reservoirs, there is still considerable important (and interesting) variation within the group (Becker et al. 2025). We also note that data collection for PREDICT was cross-sectional in nature, and therefore interpret our results as associations that warrant further mechanistic testing, for example using longitudinal or experimental approaches to test causality of interactions. Finally, the surveillance nature of the data should not be viewed as entirely representative of underlying populations. More complete sampling agnostic to host and virus taxonomy is therefore required for more confident understanding of coinfection dynamics across and within host groups. However, we note that more complete, unbiased datasets do not yet exist, and leveraging existing sampling efforts provide a foundation for future sampling priorities.

Future work could build on initial steps here by designing surveillance to more explicitly search for and report coinfection. A recent systematic review found that coinfections were common in bats, but were incidentally reported within study findings (Jones et al. 2023). More deliberate focus on sampling to capture coinfection dynamics may also require specific sampling strategies: for example, reliably detecting coinfection in an individual host requires capture of animals in-hand (versus, e.g., pooled or environmental sampling). Similarly, different parasites are associated with different organs and sites in the body, and so prioritizing particular samples or assays could be necessary, depending on study aims. Although broad sample and parasite coverage presents substantial cost considerations, several recent developments offer possible solutions. Multiplex, metagenomic, and metatranscriptomic methods all provide a fuller view of the host virome, and are especially useful when sample sizes are small due to biological limitations (e.g., collecting blood from small animals) (Boyd et al. 2015; Raghwani et al. 2023; Chang et al. 2020). Longitudinal surveillance also improves the odds of viral detection, and allows researchers to test hypotheses about asynchronous interactions between parasites, host immunity, and external environmental factors (Plowright et al. 2019). Finally, host-level data on wildlife disease are also shared infrequently (Schwantes et al. 2025), preventing the kinds of re-analyses we conducted here, or approaches that synthesize findings through systematic reviews and meta-analyses (Jones et al. 2023). Developing a better global view of wildlife disease dynamics is an essential step to conducting more macroecological research on the consequences of coinfection.

## Acknowledgements

This work was supported by NSF DBI 2515340. We thank our colleagues on the Verena team for helpful feedback and brainstorming, and the countless members of the PREDICT consortium who contributed to one of the largest and most unique ecological datasets ever collected.

## Supporting Information

**Figure S1.**
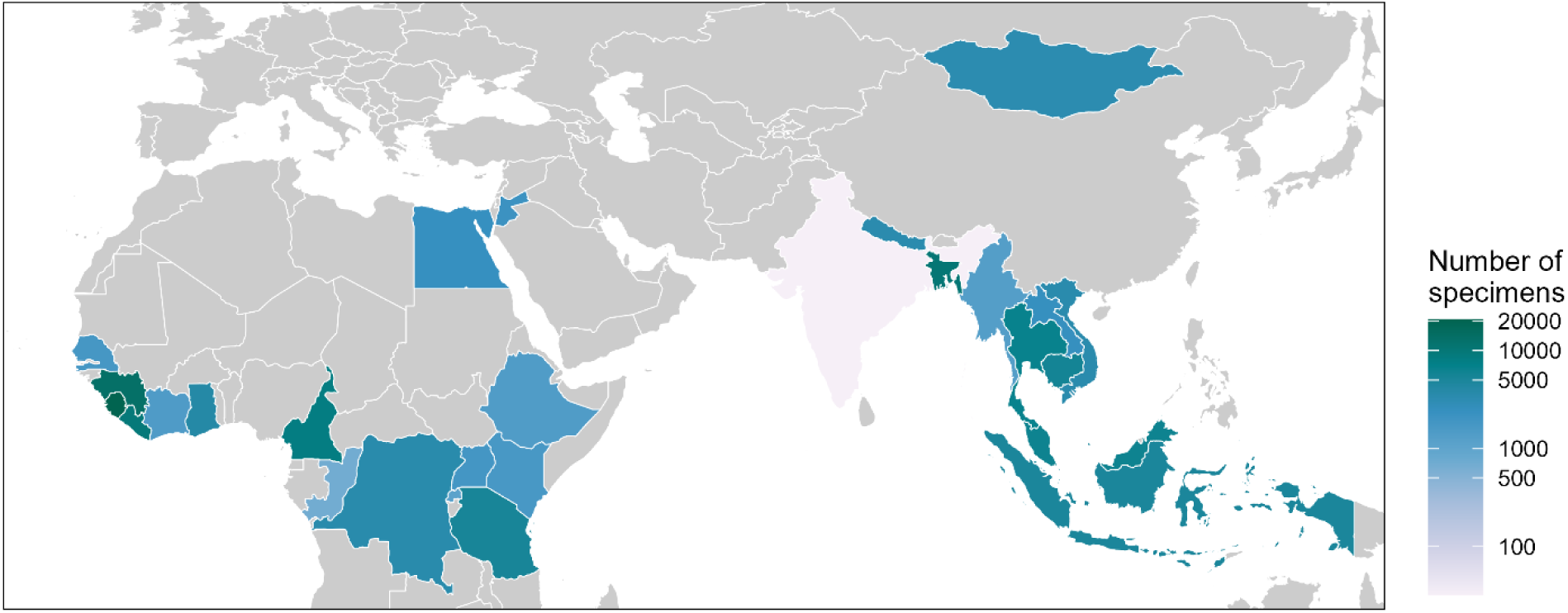
Map of sampling effort in the PREDICT-2 dataset; color corresponds to the number of unique specimens collected in a country. Note that the color legend is on a log scale.

**Figure S2.**
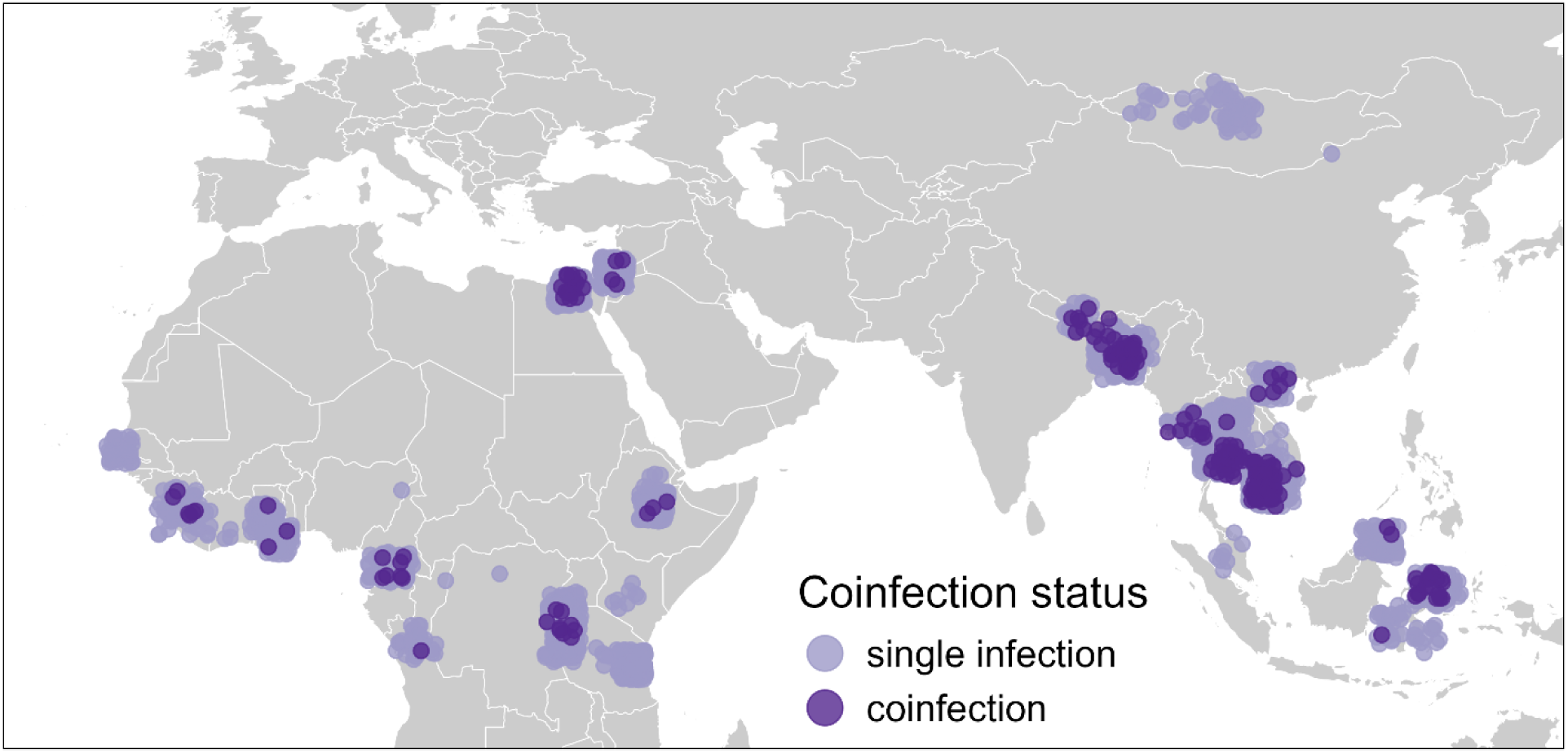
Geographic distribution of unique animals infected with one or more viruses. Points are partially transparent and jittered for improved visibility.

**Figure S3.**
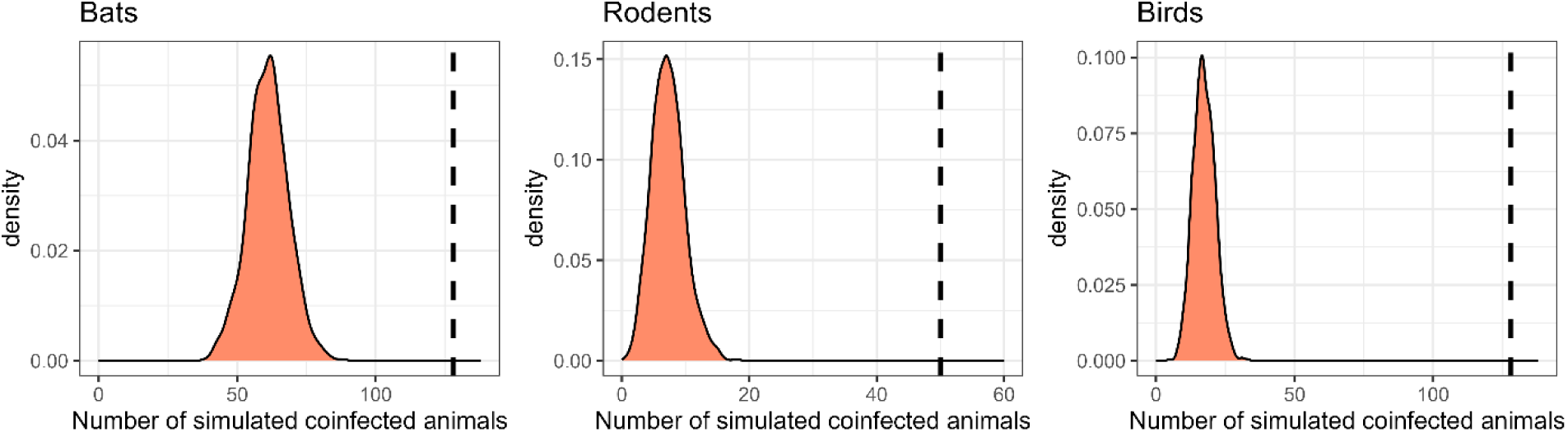
Density plots of 1000 coinfection simulations each for bats (*n* = 36,181), rodents (*n* = 12,876), and birds (*n* = 5,617). The observed number of coinfected animals (in the PREDICT-2 data) for each taxonomic group is plotted with a dashed vertical line.

**Table S1.** Number of infected animals and coinfected animals within the PREDICT-2 dataset, separated by taxonomic group. NA = not applicable (if no animals were infected, then none could be coinfected).

| **Animal taxonomic group** | **Total number tested** | **Number infected ($\geq 1$ viruses detected)** | **Number coinfecting ($\geq 2$ viruses detected)** |
| --- | --- | --- | --- |
| Bats | 36181 | 2059 | 128 |
| Birds | 5617 | 487 | 39 |
| Camels | 356 | 0 | NA |
| Carnivores | 406 | 1 | 0 |
| Cats | 62 | 0 | NA |
| Cattle/Buffalo | 517 | 2 | 0 |
| Dogs | 961 | 10 | 0 |
| Goats/Sheep | 441 | 4 | 0 |
| Horses | 1 | 0 | NA |
| Non-human primates | 6196 | 0 | NA |
| Pangolins | 332 | 0 | NA |
| Rodents | 12876 | 549 | 50 |
| Shrews | 1050 | 33 | 1 |
| Swine | 611 | 125 | 5 |
| Ungulates | 32 | 0 | NA |
| Other | 23 | 1 | 0 |

**Table S2.**
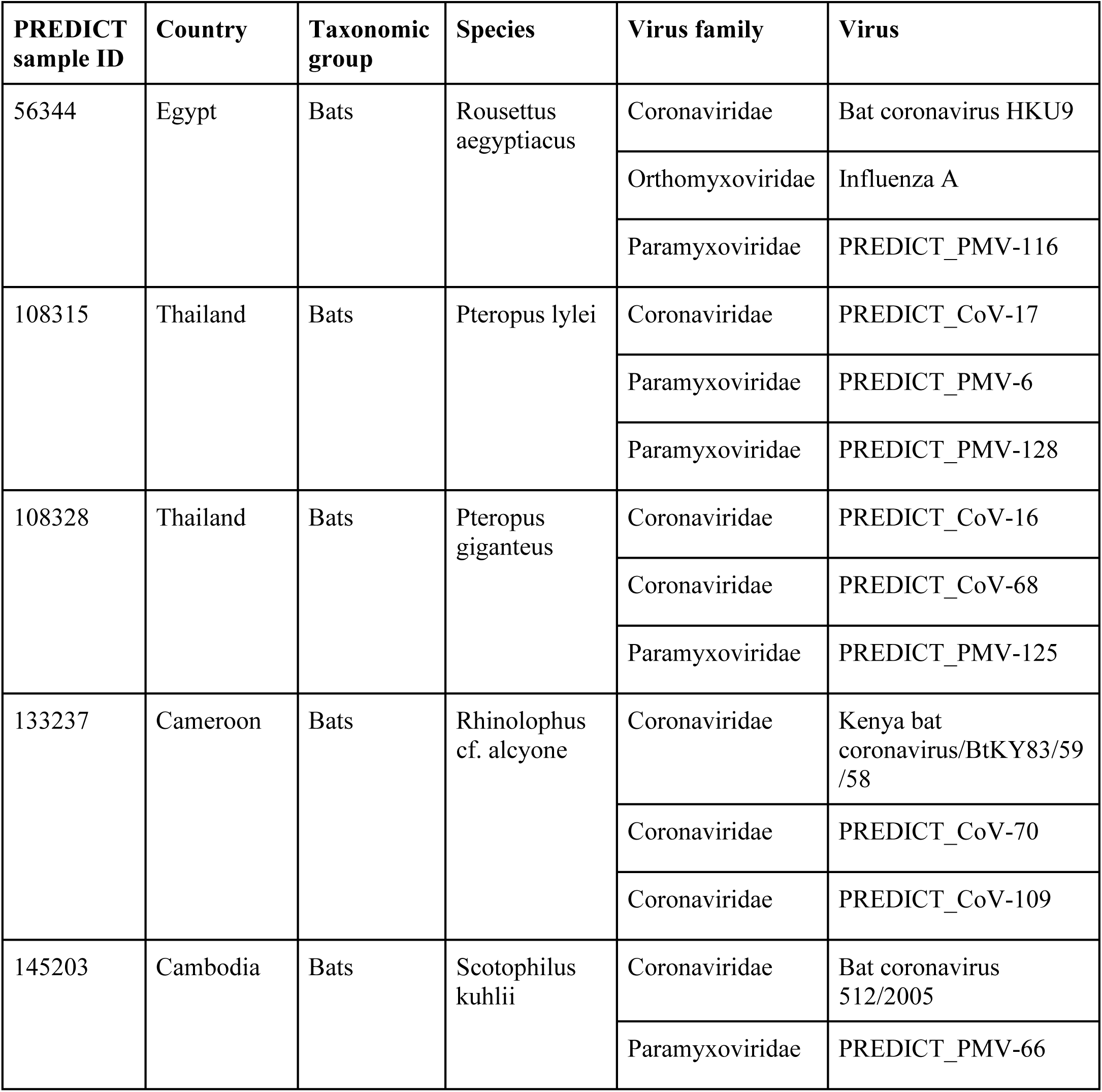
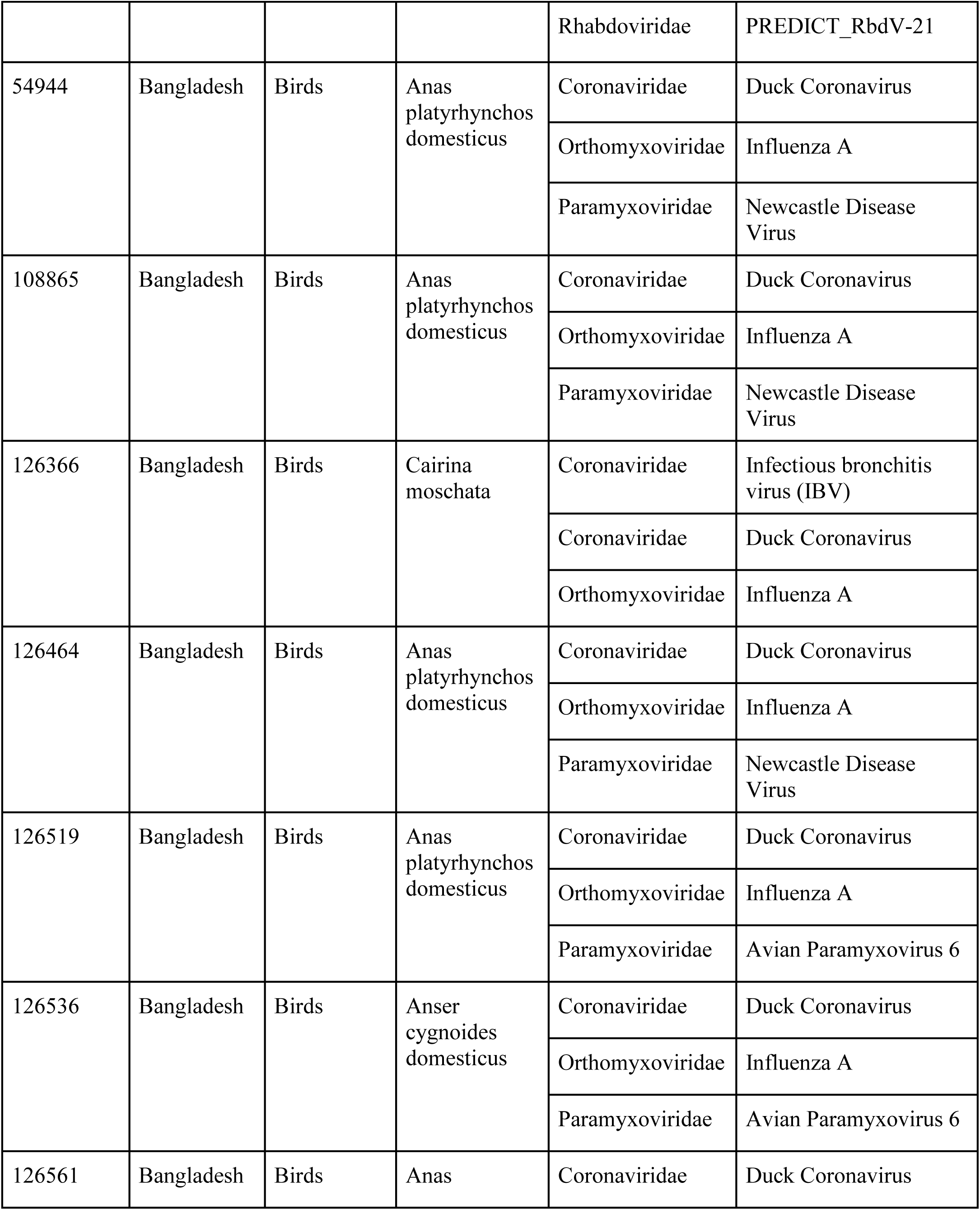
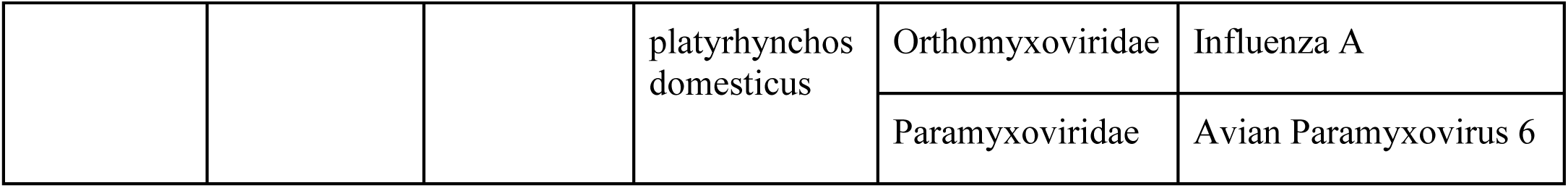
Characteristics of the 12 animals that were infected with three viruses. Note that host and virus names are provided as originally described in the PREDICT dataset, and may not conform to current taxonomies.

**Table S3.** Model coefficients for a generalized linear model to examine effects of taxonomy, sex, age, captivity status, and testing effort on virus coinfection status (binary: single infection or coinfection) in bats, birds, rodents, and shrews. *p* values < 0.05 are in bold.

| Variable |  | Estimate | SE | z | *p* | Odds ratio<br>(95% CI) |
| --- | --- | --- | --- | --- | --- | --- |
| Intercept |  | -3.51 | 0.31 | -11.20 | < **0.001** | 0.03 (0.02-0.05) |
| Geographic region | Africa and West Asia | — | — | — | — | — |
|  | South, East, and Southeast Asia | 0.88 | 0.20 | 4.35 | < **0.001** | 2.42 (1.63-3.61) |
| Taxonomic group | bats | — | — | — | — | — |
|  | birds | -1.42 | 0.45 | -3.18 | **0.001** | 0.24 (0.09-0.55) |
|  | rodents | -0.60 | 0.25 | -2.36 | 0.018 | 0.55 (0.33-0.89) |
|  | shrews | -0.90 | 1.03 | -0.88 | 0.382 | 0.41 (0.02-1.96) |
| Sex | female | — | — | — | — | — |
|  | male | -0.06 | 0.16 | -0.39 | 0.697 | 0.94 (0.68-1.29) |
| Age class | adult | — | — | — | — | — |
|  | subadult | 0.12 | 0.20 | 0.58 | 0.563 | 1.13 (0.74-1.66) |
|  | juvenile | -1.14 | 0.53 | -2.17 | **0.030** | 0.32 (0.10-0.80) |
| Captivity status | free-ranging wild animal | — | — | — | — | — |
|  | owned domesticated animal | 0.27 | 0.60 | 0.44 | 0.658 | 1.31 (0.39-4.38) |
|  | wild animal<br>in captivity | 0.75 | 0.30 | 2.48 | **0.013** | 2.12 (1.16-3.82) |
| Number of<br>virus families<br>tested |  | 0.11 | 0.05 | 2.04 | **0.042** | 1.11 (1.00-1.24) |

**Table S4.** Model coefficients (fixed effects only) for a generalized linear mixed model to examine effects of sex, age, cave roosting behavior, and testing effort on virus coinfection status (binary: single infection or coinfection) in bats. *p* values < 0.05 are in bold.

| Variable |  | Estimate | SE | z | *p* | Odds ratio (95% CI) |
| --- | --- | --- | --- | --- | --- | --- |
| Intercept |  | -2.97 | 0.64 | -4.61 | < **0.001** | 0.05 (0.01-0.18) |
| Geographic region | Africa and West Asia | — | — | — | — | — |
|  | South, East, and Southeast Asia | 1.03 | 0.23 | 4.44 | < **0.001** | 2.79 (1.78-4.39) |
| Sex | female | — | — | — | — | — |
|  | male | -0.08 | 0.21 | -0.40 | 0.686 | 0.92 (0.61-1.38) |
| Age class | adult | — | — | — | — | — |
|  | subadult | -0.53 | 0.32 | -1.64 | 0.102 | 0.59 (0.32-1.11) |
|  | juvenile | -2.03 | 0.73 | -2.77 | **0.006** | 0.13 (0.03-0.55) |
| Cave roosting | no | — | — | — | — | — |
|  | yes | -0.07 | 0.29 | -0.26 | 0.796 | 0.93 (0.53-1.63) |
| Number of virus families tested |  | -0.03 | 0.09 | -0.33 | 0.741 | 0.97 (0.81-1.16) |

**Table S5.** Model coefficients for a generalized linear model to examine effects of sex, age, and testing effort on virus coinfection status (binary: single infection or coinfection) in rodents. *p* values < 0.05 are in bold.

| Variable |  | Estimate | SE | z | *p* | Odds ratio (95% CI) |
| --- | --- | --- | --- | --- | --- | --- |
| Intercept |  | -5.79 | 0.86 | -6.76 | < **0.001** | 0.00 (0.00-0.01) |
| Sex | female | — | — | — |  | — |
|  | male | 0.13 | 0.32 | 0.42 | 0.676 | 1.14 (0.61-2.17) |
| Age class | adult | — | — | — |  | — |
|  | subadult/juvenile | 0.61 | 0.33 | 1.84 | 0.066 | 1.84 (0.95-3.51) |
| Number of virus families tested |  | 0.45 | 0.10 | 4.36 | < **0.001** | 1.57 (1.29-1.94) |

**Table S6.** Expected versus observed coinfection prevalence for virus families, for all animals. *P* values were calculated from binomial tests and adjusted using a Bonferroni correction. *P_adj_* values < 0.05 are in bold.

| **Virus family combination** | **Expected coinfection prevalence** | **Observed coinfection prevalence** | **Observed > expected** | ***P<sub>adj</sub>*** |
| --- | --- | --- | --- | --- |
| Arenaviridae-Arenaviridae | 5.80e-9 | 0 | No | 1 |
| Arenaviridae-Coronaviridae | 2.76e-6 | 0 | No | 1 |
| Arenaviridae-Filoviridae | 1.39e-8 | 0 | No | 1 |
| Arenaviridae-Flaviviridae | 5.80e-9 | 0 | No | 1 |
| Arenaviridae-Hantaviridae | 4.64e-9 | 0 | No | 1 |
| Arenaviridae-Orthomyxoviridae | 5.49e-7 | 0 | No | 1 |
| Arenaviridae-Paramyxoviridae | 5.42e-7 | 0 | No | 1 |
| Arenaviridae-Rhabdoviridae | 5.68e-8 | 0 | No | 1 |
| Coronaviridae-Coronaviridae | 1.31e-3 | 1.64-3 | Yes | 0.841 |
| Coronaviridae-Filoviridae | 6.62e-6 | 0 | No | 1 |
| Coronaviridae-Flaviviridae | 2.76e-6 | 0 | No | 1 |
| Coronaviridae-Hantaviridae | 2.21e-6 | 0 | No | 1 |
| Coronaviridae-Orthomyxoviridae | 2.61e-4 | 6.85e-4 | Yes | **< 0.001** |
| Coronaviridae-Paramyxoviridae | 2.58e-4 | 1.05e-3 | Yes | **< 0.001** |
| Coronaviridae-Rhabdoviridae | 2.70e-5 | 4.57e-5 | Yes | 1 |
| Filoviridae-Filoviridae | 3.34-8 | 0 | No | 1 |
| Filoviridae-Flaviviridae | 1.39e-8 | 0 | No | 1 |
| Filoviridae-Hantaviridae | 1.11e-8 | 0 | No | 1 |
| Filoviridae-Orthomyxoviridae | 1.32e-6 | 0 | No | 1 |
| Filoviridae-Paramyxoviridae | 1.30e-6 | 0 | No | 1 |
| Filoviridae-Rhabdoviridae | 1.36e-7 | 0 | No | 1 |
| Flaviviridae-Flaviviridae | 5.80e-9 | 0 | No | 1 |
| Flaviviridae-Hantaviridae | 4.64e-9 | 0 | No | 1 |
| Flaviviridae-Orthomyxoviridae | 5.49e-7 | 1.52e-5 | Yes | 1 |
| Flaviviridae-Paramyxoviridae | 5.42e-7 | 0 | No | 1 |
| Flaviviridae-Rhabdoviridae | 5.68e-8 | 0 | No | 1 |
| Hantaviridae-Hantaviridae | 3.71e-9 | 0 | No | 1 |
| Hantaviridae-Orthomyxoviridae | 4.39e-7 | 0 | No | 1 |
| Hantaviridae-Paramyxoviridae | 4.33e-7 | 0 | No | 1 |
| Hantaviridae-Rhabdoviridae | 4.55e-8 | 0 | No | 1 |
| Orthomyxoviridae-Orthomyxoviridae | 5.19e-5 | 0 | No | 1 |
| Orthomyxoviridae-Paramyxoviridae | 5.12e-5 | 1.52e-4 | Yes | 0.090 |
| Orthomyxoviridae-Rhabdoviridae | 5.38e-6 | 0 | No | 1 |
| Paramyxoviridae-Paramyxoviridae | 5.06e-5 | 7.61e-5 | Yes | 1 |
| Paramyxoviridae-Rhabdoviridae | 5.31e-6 | 9.14e-5 | Yes | **< 0.001** |
| Rhabdoviridae-Rhabdoviridae | 5.57e-7 | 0 | No | 1 |

**Table S7.**
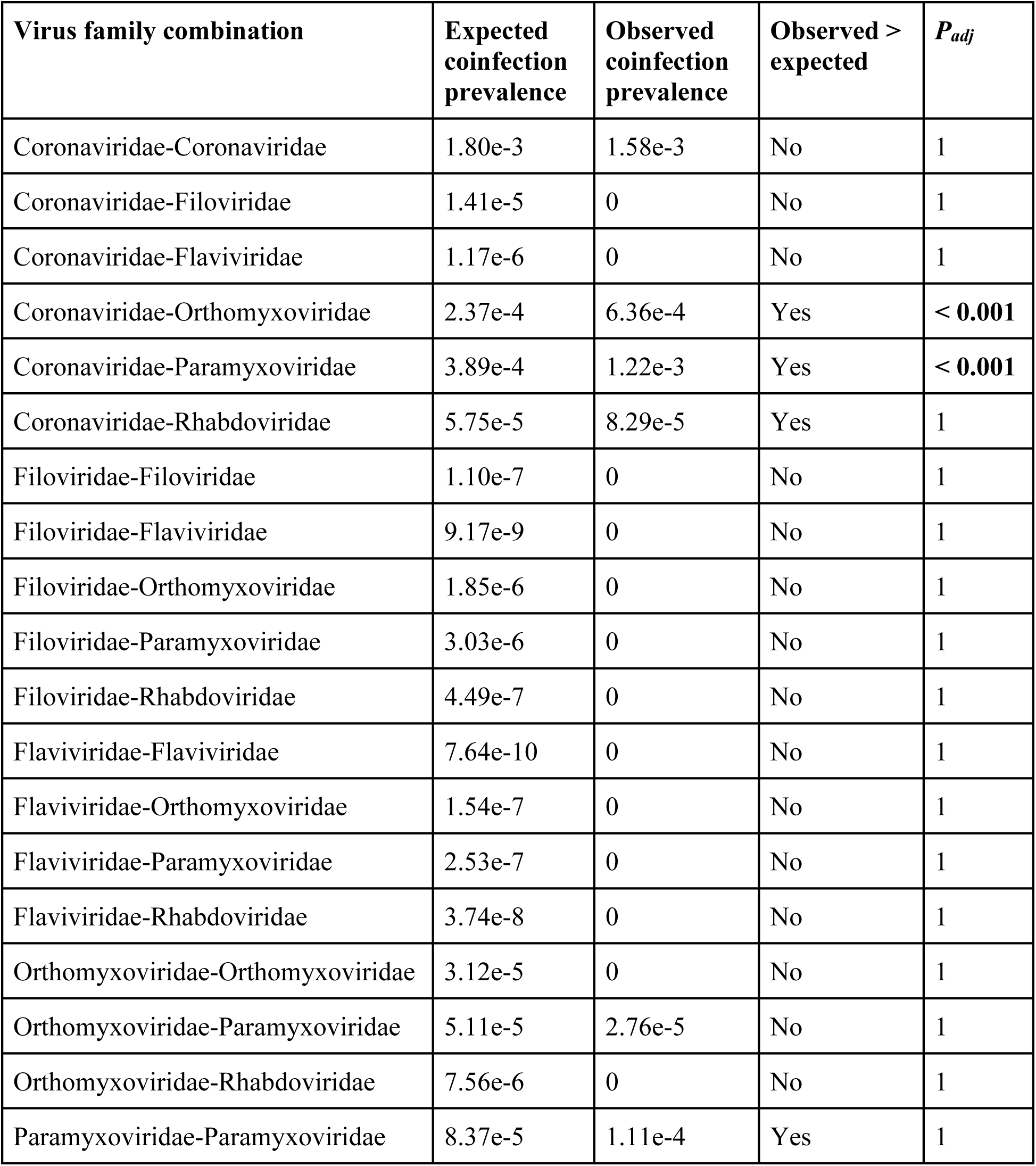
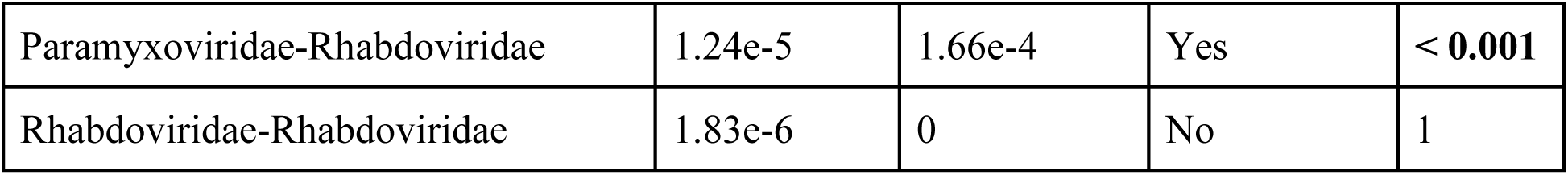
Expected versus observed coinfection prevalence for virus families, for bats only. *P* values were calculated from binomial tests and adjusted using a Bonferroni correction. *P_adj_* values < 0.05 are in bold.

**Table S8.** Expected versus observed coinfection prevalence for virus families, for rodents only. *P* values were calculated from binomial tests and adjusted using a Bonferroni correction. *P_adj_* values < 0.05 are in bold.

| **Virus family combination** | **Expected coinfection prevalence** | **Observed coinfection prevalence** | **Observed > expected** | **$P_{adj}$** |
| --- | --- | --- | --- | --- |
| Arenaviridae-Arenaviridae | 1.51e-7 | 0 | No | 1 |
| Arenaviridae-Coronaviridae | 1.47e-5 | 0 | No | 1 |
| Arenaviridae-Hantaviridae | 6.03e-8 | 0 | No | 1 |
| Arenaviridae-Orthomyxoviridae | 6.03e-8 | 0 | No | 1 |
| Arenaviridae-Paramyxoviridae | 1.60e-6 | 0 | No | 1 |
| Coronaviridae-Coronaviridae | 1.44e-3 | 3.73e-3 | Yes | **< 0.001** |
| Coronaviridae-Hantaviridae | 5.90e-6 | 0 | No | 1 |
| Coronaviridae-Orthomyxoviridae | 5.90e-6 | 0 | No | 1 |
| Coronaviridae-Paramyxoviridae | 1.56e-4 | 1.55e-4 | No | 1 |
| Hantaviridae-Hantaviridae | 2.41e-8 | 0 | No | 1 |
| Hantaviridae-Orthomyxoviridae | 2.41e-8 | 0 | No | 1 |
| Hantaviridae-Paramyxoviridae | 6.39e-7 | 0 | No | 1 |
| Orthomyxoviridae-Orthomyxoviridae | 2.41e-8 | 0 | No | 1 |
| Orthomyxoviridae-Paramyxoviridae | 6.39e-7 | 0 | No | 1 |
| Paramyxoviridae-Paramyxoviridae | 1.69e-5 | 0 | No | 1 |

